# Effect of cryopreservation and post-cryopreservation somatic embryogenesis on the epigenetic fidelity of cocoa (*Theobroma cacao* L.)

**DOI:** 10.1101/057273

**Authors:** Raphael Adu-Gyamfi, Andy Wetten, Carlos Marcelino, Rodríguez López

## Abstract

While cocoa plants regenerated from cryopreserved somatic embryos can demonstrate high levels of phenotypic variability, little is known about the sources of the observed variability. Previous studies have shown that the encapsulation-dehydration cryopreservation methodology imposes no significant extra mutational load since embryos carrying high levels of genetic variability are selected against during protracted culture. Also, the use of secondary rather than primary somatic embryos has been shown to further reduce the incidence of genetic somaclonal variation. Here, the effect of *in vitro* conservation, cryopreservation and post-cryopreservation generation of somatic embryos on the appearance of epigenetic somaclonal variation were comparatively assessed. To achieve this we compared the epigenetic profiles, generated using Methylation Sensitive Amplified Polymorphisms, of leaves collected from the ortet tree and from cocoa somatic embryos derived from three *in vitro* conditions: somatic embryos, somatic embryos cryopreserved in liquid nitrogen and somatic embryos generated from cryoproserved somatic embryos. Somatic embryos accumulated epigenetic changes but these were less extensive than in those regenerated after storage in LN. Furthermore, the passage of cryopreserved embryos through another embryogenic stage led to further increase in variation. Interestingly, this detected variability appears to be in some measure reversible. The outcome of this study indicates that the cryopreservation induced phenotypic variability could be, at least partially, due to DNA methylation changes.

**Key message:** Phenotypic variability observed in cryostored cocoa somatic-embryos is epigenetic in nature. This variability is partially reversible, not stochastic in nature but a directed response to the *in-vitro* culture and cryopreservation.

## Introduction

The propagation of plant material through *in vitro* culture can lead to marked increases in the frequency of variants [1]. These enhanced frequencies can be higher than those associated with the use of mutagens [2] and have been termed somaclonal variation [3]. The nature of such variation can be genetic (altering the DNA sequence of the ramets) and/or epigenetic (which does not affect the DNA sequence but affects its chemistry and structure, gene expression and may ultimately induce some form of phenotypic abnormality [4]. The addition of a methyl group to cytosine residues (cytosine methylation) is probably the most studied feature of epigenetic regulation in Eukaryotes. In plants, it occurs in three sequence contexts: CG, CNG, or CNN (in order of relative abundance) (N = any nucleotide other than G [5]). Cytosine methylation occurring within promoters or coding regions typically acts to repress gene transcription by changing local chromatin structure [6], thereby preventing the binding of DNA-binding proteins to the promoter regions [7] or as a binding cue for transcriptional repressions [8].

Different factors have been reported to affect the level of both genetic and epigenetic somaclonal variation induced during *in vitro* culture, including: **1.** Medium composition [9,10]. **2.** The propagation technique used, with methods involving a dedifferentiated callus phase expected to show higher levels of variability [11]. **3.** Origin of the donor tissue, with regenerants maintaining epigenetic features of the explant tissue used for propagation [12]. **4.** Time in culture and number of regeneration events [12]. Somaclonal variation generally increases with prolonged callus phase and with the number of multiplication cycles [3,13]. However, Rodriguez Lopez et al. [12] observed that older cocoa calli, as well as exhibiting reduced embryogenic potential, yielded somatic embryos (SE) containing less genetic and epigenetic aberrations. They hypothesised that totipotent cell lineages with few or no mutations are selected during protracted cultures. In cocoa, primary somatic embryogenesis has previously been shown to induce high levels of chimeric [14] and homogenous [12,15] genetic and epigenetic mutants. Nevertheless, in the same studies fewer genetic variants were detected among secondary somatic embryos (SSEs) than among primary somatic embryos (PSEs). These findings are in line with Fang et al. [16] who reported an absence of mutations in the SSE population of cocoa studied probably due to SSEs being derived directly from epidermal cells rather than from dedifferentiated callus.

Due to the difficulty in generating large quantities of cocoa plants via cuttings and to the recalcitrant nature of cocoa seed with regard to low temperature storage, cryopreservation of tissue-cultured germplasm represents the most attractive backup for vulnerable field collections of the species. The lengthy culture periods associated with the establishment of embryogenic cocoa callus lines also mean that it will be a wise precaution to back up large *in vitro* germplasm collections with cryopreserved cell lines. Different groups have shown that the encapsulation-dehydration [16] and cryopreservation of plant material [17] do not impose a significant extra genetic mutational load to the plants regenerated *in vitro*. However, little is known about how the cryopreservation methodology and subsequent propagation cycles affect the fidelity of the regenerant plants from an epigenetic perspective. Methylation-sensitive amplified polymorphism (MSAP) is a modification of Amplified Fragment Length Polymorphism that makes use of differential sensitivity of certain restriction endonucleases to cytosine methylation to study the level and global patterns of methylation across the target genomes [18]. MSAP employs a pair of isoschizomer enzymes (*Hpa*II and *MspI*) with differential sensitivity to methylation in their recognition sequence. In short, *Hpa*II is inactive (does not cut its recognition site, 5’-CCGG-3’) if one or both cytosines are methylated on both DNA strands, but cleaves when all cytosines are unmethylated or if one or both cytosines are methylated in only one strand. MspI is, in contrast, considered by convention methylation to be insensitive and cleaves if all cytosines are unmethylated, and if the internal cytosine is methylated but not when the external cytosine is methylated. MSAP analysis has been previously used for the detection of tissue culture induced epigenetic changes in a large number of plant taxa [19], including important crops such as: tobacco [9], rice [20], oil palm [21], barley [22], pea [23], potato [24], cocoa [12], grapevine [25], cassava [26], and garlic [27]. In this study we used the MSAP technology to determine the effect of cryopreservation and post-cryopreservation regeneration of tertiary SEs (TSEs) on the appearance of somaclonal variation compared to equivalent variation in long term *in vitro* conservation of cocoa SSEs.

## Materials and Methods

### Plant Material

All the SEs used in this study were generated from a single AMAZ 15 cocoa tree held in the International Cocoa Quarantine Centre, Reading, UK, with somatic embryogenesis initiated from staminode cultures as described in [28]. Three groups of propagules plus a set of reference samples from the ortet tree were compared: **1** SSEs maintained in ED medium [28] culture for the 357 d duration of the experiment (*‘in vitro’* hereafter); **2** SSEs recovered after 1 h storage in liquid nitrogen (LN) using vitrification-based cryopreservation [29] (‘1 h LN’ hereafter); **3** TSEs regenerated from 1 h LN cotyledon explants following [28] (‘post 1 h LN’ hereafter) and **4** the ortet tree reference set were 6 mm discs obtained from 24 randomly selected recently expanded leaves from the donor tree. All samples were kept at −20°C until DNA extraction (see Fig 1 and Table 1).

**Fig 1.**
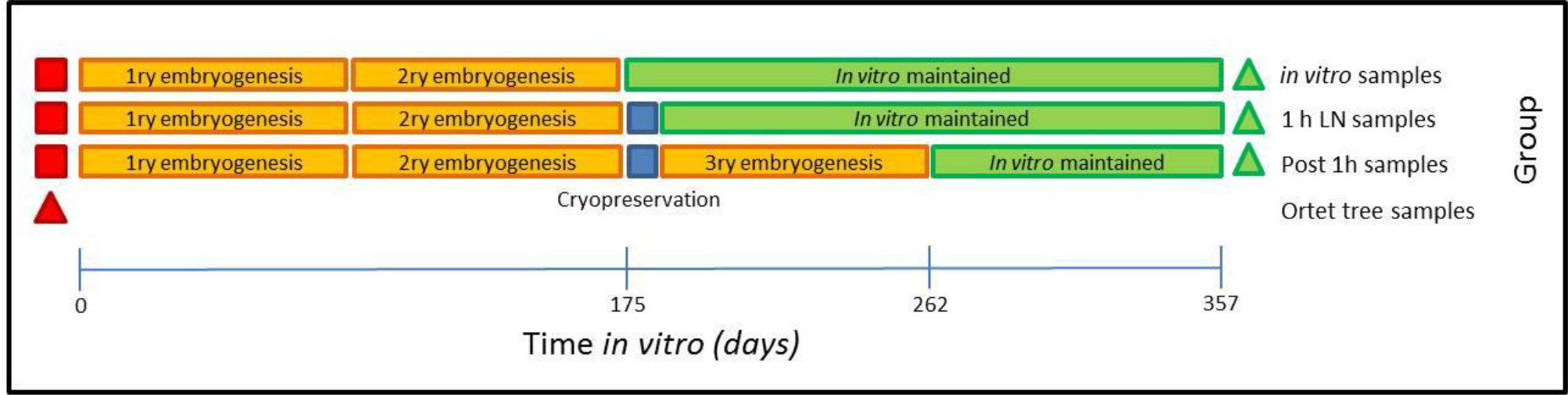
Schematic representation of experimental design. Somatic embryogenesis was initiated from staminodes (red squares) obtained from a single AMAZ 15 cocoa tree from the International Cocoa Quarantine Centre, Reading, UK. Triangles represent 24 samples collected from individual newly expanded leaves for DNA extraction from each group (Donor plant (Red) and somatic embryo derived plants (Green)). Orange boxes represent successive somatic embryogenic events (as described in [28]). Blue boxes represent a cryopreservation event (1 h in liquid nitrogen) of somatic embryos (as described [29]). Green boxes represent maintenance of somatic embryos *in vitro* on ED medium [28]. Horizontal bar represents accumulated time spend under *in vitro* conditions by the samples used in this study

**Table 1.**
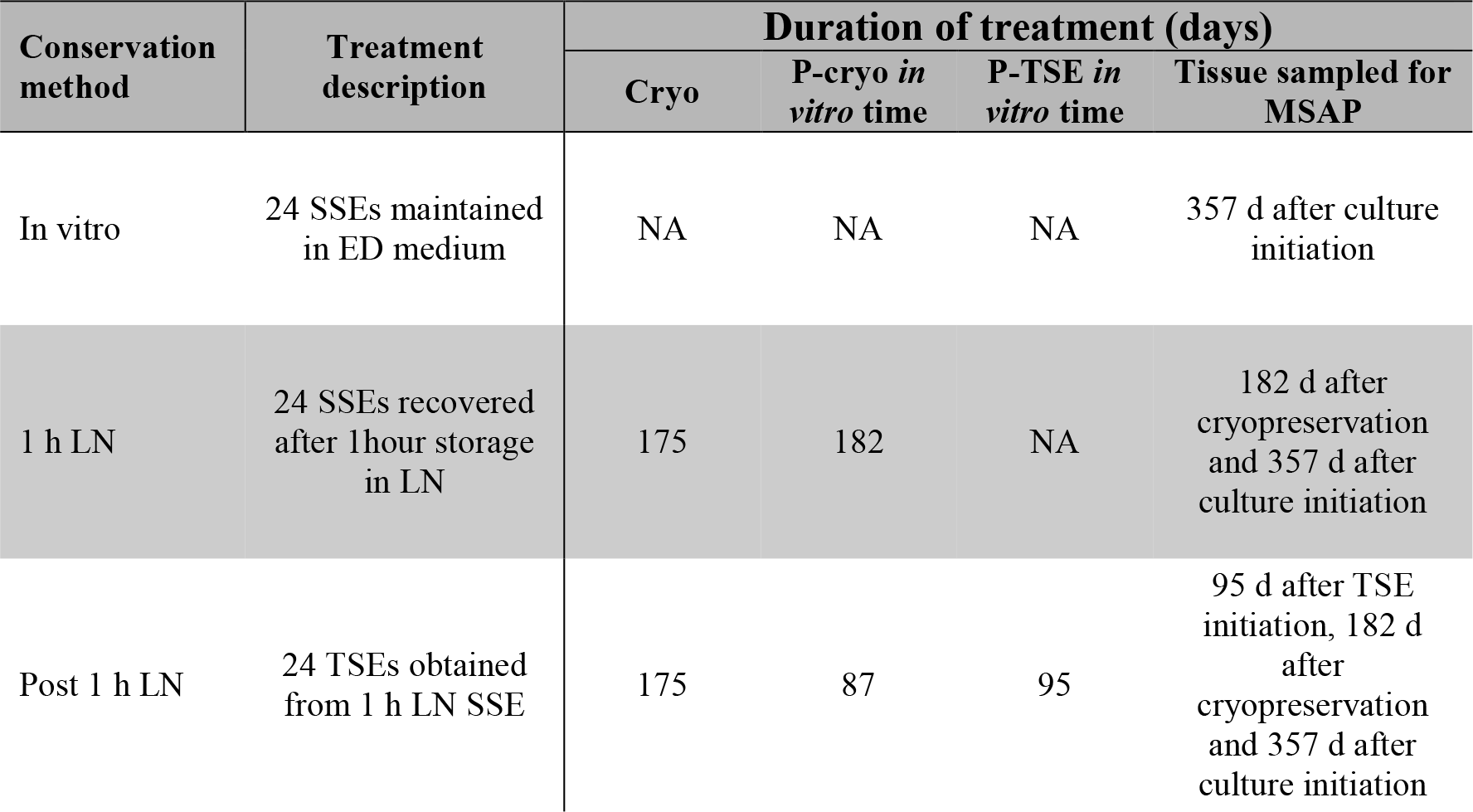
Description of cocoa samples used for MSAP analysis.

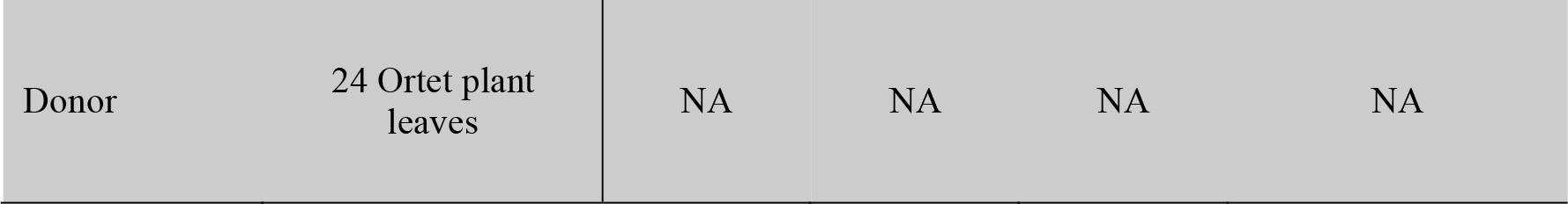

**NA**: not applicable. **Cryo:** Number of days since the initiation of somatic embryos from staminodes, prior to samples being subjected to cryopreservation. **P-cryo *in vitro* time:** Number of days that secondary somatic embryos (SSEs) spent in ED medium [28] since they were removed from cryopreservation. **P-TSE** *in vitro* **time:** Number of days that tertiary somatic embryos (TSEs) spent in ED medium [28] since the tertiary somatic embryogenesis event.

### DNA extraction

Genomic DNA was extracted from leaves of somatic embryos and donor plant using the DNeasy 96 plant kit (Qiagen, UK) following manufacturer’s instructions and eluted in 50 μl Buffer AE (Qiagen). Two replicate extractions were performed for each sample. The DNA was quantified on a NanoDrop^TM^ 2000c (ThermoLifescience). Extracted DNAs were diluted with nanopure water to produce working stocks of 10 ng/μl.

### MSAP procedure

A modification of the original MSAP method, as described by Rodriguez Lopez et al. [30], was used on all DNA extractions. In short, the method consists of the parallel digestion of genomic DNA with two methylation-sensitive isoschizomers (*MspI* and *HpaII*) as frequent cutters, each in combination with the same rare cutter (*EcoRI*), adaptor ligation, followed by two selective PCR amplifications with primers complementary to the adaptors but with unique 3’ overhangs (Table 2). Thirty-two primer combinations consisting of eight +3EcoRI and four +*2HpaII/MspI* combinations were evaluated with a subset of 8 randomly selected pre-selective amplifications containing two samples from each group described in Table 1. The evaluation was done to assess the level of intra-treatment variation for each primer combination and their ability to generate informative and consistent MSAP profiles. Based on these results the best two primer combinations (*Hp1/Eco1* and *Hp3/Eco3*) were designated as primers for the selective amplification of 96 samples from the four experimental groups. Resultant PCR products labelled with FAM fluorescent dye were diluted 1/10 in sterile nano water and 1 μl was combined with 1 μl ROX/HiDi mix (50 μl ROX plus 1 ml HiDi formamide). Samples were then denatured by heating at 95 °C for 5 min, snap-cooled on ice for 2 min and run on an ABI PRISM 3100 Genetic Analyzer Capillary Sequencer (16 capillary array model) at 3 kV for 22 s and 15 kV for 45 min at 60 °C.

**Table 2.**
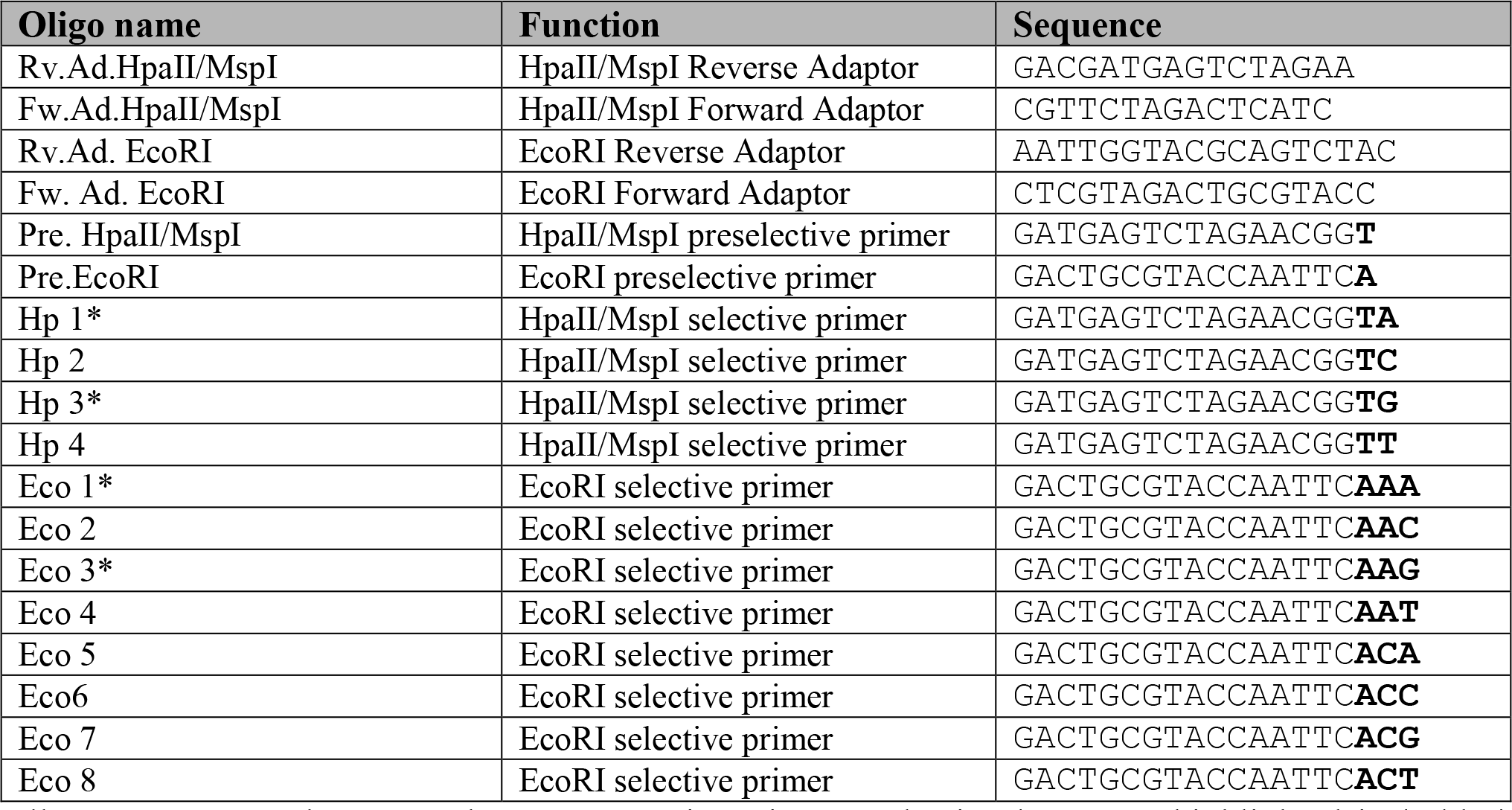
Sequence of oligonucleotides used for MSAP analysis.

All sequences are shown on the 5’ to 3’ orientation. 3’selective bases are highlighted in bold. * indicates primers chosen for selective amplification.

### Data analysis

GeneMapper_®_ Software Version 4.0 was used to generate a binary data matrix of bands present (1) or absent (0) on the MSAP profiles from samples restricted using *EcoRI* and *HpaII* or *MspI* and amplified using the different primer combinations shown in Table 2. Only epiloci ranging from 100 to 500bp in size were considered for analysis. Fragments present/absent in all but one individual were considered uninformative and removed from all data sets. GenAlex [31] 6.5 software was used to analyse the binary data matrix as described in Rodriguez Lopez et al. [30]. In brief, first the frequency of all epiloci was calculated for each group. Epiloci were considered unique to *in vitro* culture samples (*in vitro*, 1 h LN and/or Post 1 h LN) when the their frequency was higher than 75% in the donor plant samples and lower than 25% in the *in vitro* samples and *vice versa*. Equally, epiloci were considered unique to cryopreserved samples (1 h LN and/or Post 1 h LN) when their frequency was higher than 75% in the 1h LN samples and lower than 25% in the *in vitro* samples and *vice versa.* Finally, epiloci were considered associated to *in vitro* culture or cryopreservation when their frequencies where higher than 60% in the donor plant/1h LN samples and lower than 30% in the *in vitro* samples and *vice versa.* Next using GenAlex (v.6.4), the epigenetic variability between ortets (samples from donor plant) and ramet groups (samples generated under different *in vitro* conditions, Table 1) was visualized by Principal Coordinate Analysis (PCoA) based on the amalgamation of the MSAP profiles obtained from samples restricted with *HpaII* or *MspI* and amplified with primer combinations *Hp1/Eco1* and *Hp3/Eco3.* Analysis of Molecular Variance (AMOVA) was then used to estimate and test the significance of the epigenetic diversity within and between the different groups. Pairwise PhiPT comparisons between each group based on 10000 permutations was used to infer the overall level of divergence in DNA methylation between groups. Finally, the level of epigenetic variability introduced by each of the treatments was estimated using two values from the AMOVA: **1** calculated PhiPT between ortet tree samples and samples from each treatment (*in vitro*, 1h LN, Post 1h LN) (i.e., the lower the PhiPT, the lower the level of DNA methylation variability associated by that treatment), **2** The mean sum of squares within population (SSWP) was used to infer epigenetic variation within treatments (the higher the mean SSWP, the higher the level of epigenetic variability associated by that treatment) [32].

## Results

### Effect of conservation methods on somatic embryo methylation patterns

To analyse the effect of the different *in vitro* conservation methods on the DNA methylation patterns of cocoa SE we compared the MSAP profiles of samples from the donor plant (AMAZ 15) to those obtained from SE conserved as described in Table 1. A total of 220 fragments ranging from 100bp to 500bp were generated by primer combinations *Hp1/Eco1* and *Hp3/Eco3* (108 and 112 fragments respectively). Of these, 3.6% (8 epiloci) and 8.6% (19 epiloci) were deemed unique or induced by cryopreservation and *in vitro* culture respectively (Table 3). In total 27 *in vitro* cultures and cryopreservation markers were obtained, 13 using *HpaII* and 14 using MspI (Table 3). Of these markers, the frequency of epialleles *HpaII* 1.1 (232), *HpaII* 1.1 (233), *HpaII* 3.3 (168),*MspI* 3.3 (303) and MspI 3.3 (407) showed a correlation with the accumulation of *in vitro* procedures (i.e. number of somatic embryogenesis and cryopreservation events) (Table 3, S1 Fig A–E). In contrast, epialleles *HpaII* 1.1 (367), *HpaII* 3.3 (181), *HpaII* 3.3 (142), *MspI* 3.3 (202) and *MspI* 1.1 (278) showed a change in frequency associated with somatic embryogenesis/cryopreservation followed by a reversion to donor tree allele frequency levels after cryopreservation or after tertiary embryogenesis (Table 3 and S1 Fig F–J).

**Table 3.**
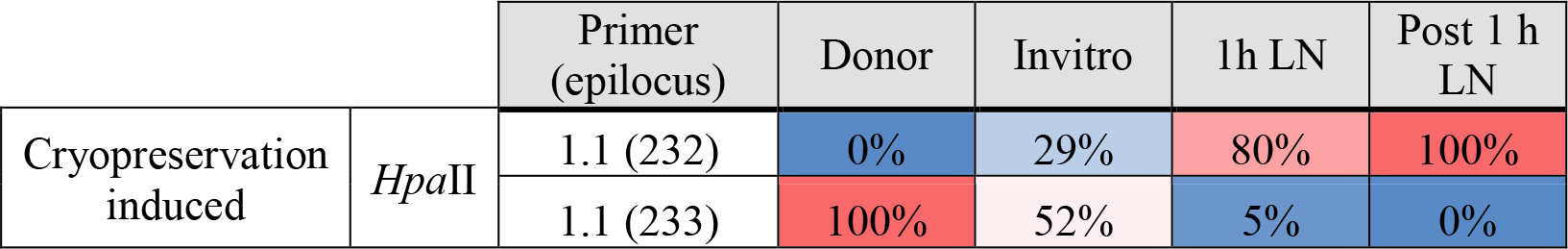
Somatic embryogenesis/cryopreservation induced epigenetic polymorphisms.

**Table 3.**
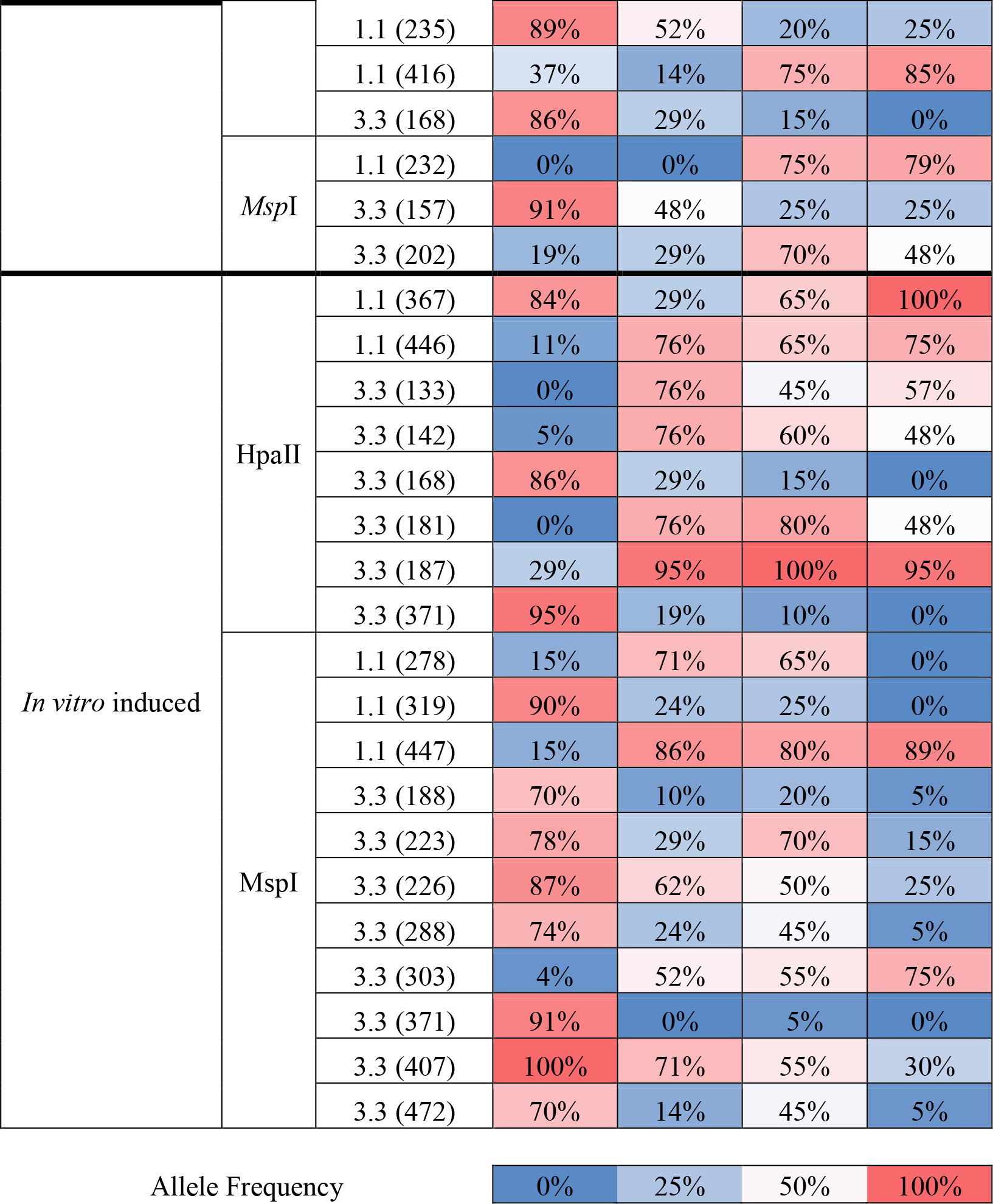

MSAP generated epiloci unique to *in vitro* culture samples (*in vitro*, 1 h LN and/or Post 1 h LN) and to cryopreserved samples (1 h LN and/or Post 1 h LN). MSAP profiles were obtained using methylation sensitive isoschizomers *Hpa*II and *Msp*I and primer combinations and *Hp1/Eco1* and *Hp3/Eco3.* Column Primer (epilocus) shows primer combination used to generate the specific MSAP product (i.e. 1.1 = *Hp1/Eco1* and 3.3 = *Hp3/Eco3*). Columns Donor, Invitro, 1 h LN and Post 1 h LN show the frequency of each epiloci in each of the compared groups (i.e. **Donor:** AMAZ 15 cocoa ortet tree used to regenerate all somatic embryos; **Invitro**: *In vitro* maintained secondary somatic embryos; **1 h LN**: Secondary somatic embryos recovered from after 1 hour storage in liquid nitrogen and **Post 1 h LN**: tertiary somatic embryos generated from 1 h LN samples).

AMOVA of profiles generated using *Hpa*II showed that 19% of the variation was explained by differences between groups while 81% was due to individual differences. Similarly, 21% of the variability detected by *Msp*I was explained by differences between groups and 79% was due to individual differences. Principal Coordinate Euclidean Analysis (PCoA) was used to provide an overview of the epigenetic variability introduced by somatic embryogenesis and cryopreservation. Overall, profiles generated using both *Msp*I and *Hpa*II provided a clear separation between donor plant samples and all SE samples (Fig 2A). However, the separation observed between the different ramet groups was slightly clearer when using profiles from samples restricted with MspI (Fig 2B) than with *Hpa*II (Figure2C). PhiPT values confirmed the epigenetic variation observed between groups using PCoA analysis, with groups presenting higher levels of epigenetic divergence when MspI profiles were used (Table 4). The largest differences between groups were detected between donor plant samples and cryopreserved samples (i.e. 1h LN and Post-1h LN) whilst the lowest levels of epigenetic divergence between groups were detected between both cryopreserved samples (i.e. 1h LN and Post-1h LN) restricted using *HpaII.* AMOVA analysis using 10000 permutations showed that all pairwise distances were significant (P>0.0002). When PhiPT values generated from *HpaII* and *MspI* MSAP profiles were taken together, Post-1h LN samples were the most epigenetically divergent from the donor samples (Fig 3A). Analysis of epigenetic variance within groups estimated using the mean SSWP showed similar levels of variability amongst donor, *in vitro* and Post-1h LN. However, higher levels of variability were detected between 1h LN samples (mean SSWP*_MspI_* = 278) compared to both donor plant samples (mean SSWP*_Mspi_* = 237) and to non-cryopreserved somatic embryos (mean SSWP*_MspI_* = 229) (Fig 3B).

**Fig 2.**
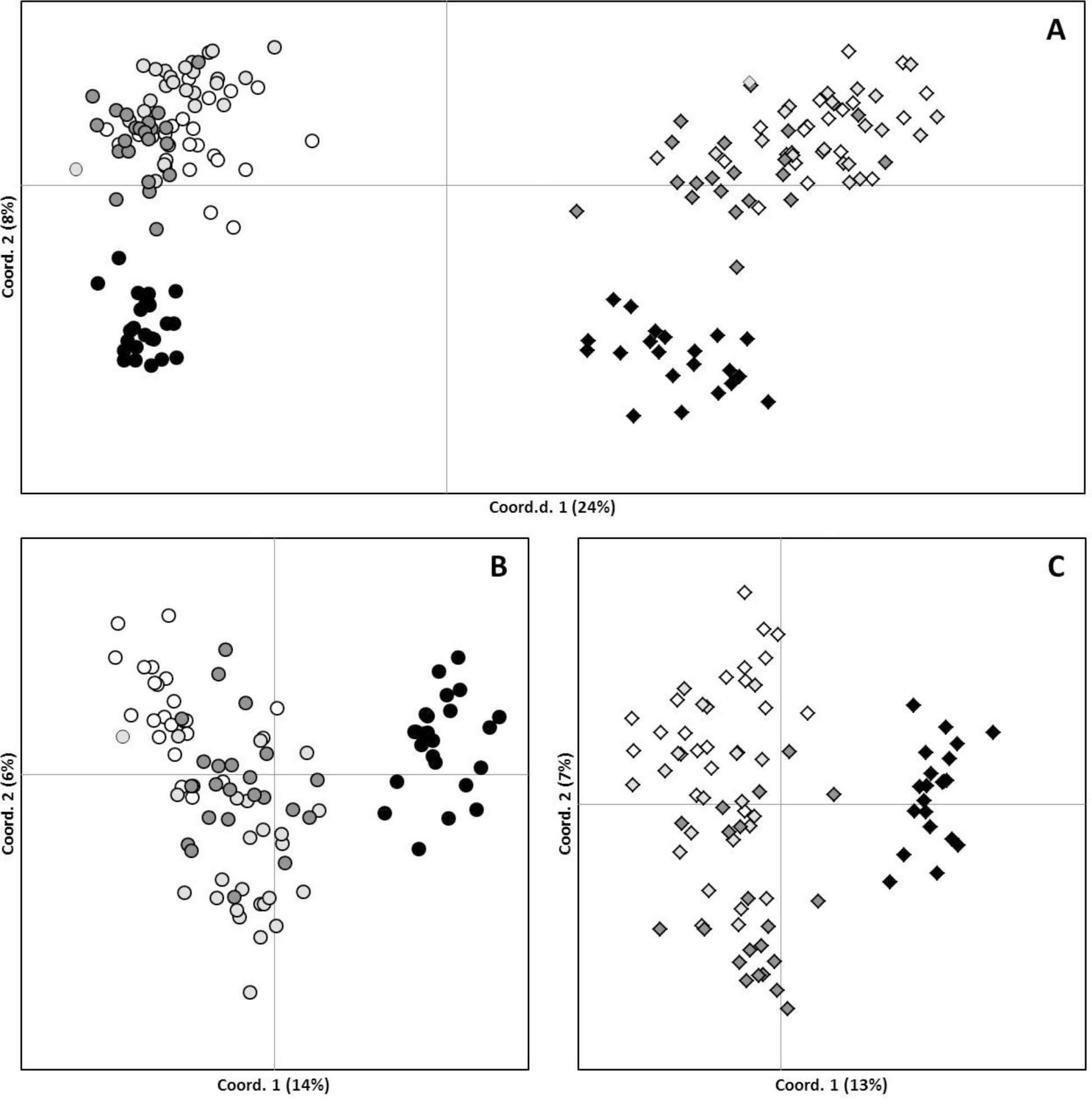
Effect of conservation method on somaclonal variation pattern in cocoa somatic embryos. Principal coordinate analysis based on Euclidean Analysis of MSAP distances between 72 somatic embryos grouped by method of conservation and 24 leaf samples of the donor tree. MSAP profiles were obtained using methylation sensitive isoschizimers MspI (circles) and *HpaII* (rhomboids) and methylation insensitive enzyme *EcoRI* primer combinations and *Hp1/Eco1* and *Hp3/Eco3*. Individual figures show PCoA analysis from MSAP profiles obtained using **(A)** both MspI and *HpaII,* **(B)** MspI only and **(C)** *HpaII* only. 1 h: SEs recovered from secondary SE after 1 hour storage in liquid nitrogen; Post 1 h: tertiary SEs generated from 1 h samples; Invitro: *In vitro* maintained secondary SEs, and Donor: AMAZ 15 cocoa ortet tree used to regenerate SEs. Hp or Msp preceding a treatment means MSAP profiles were generated using restriction enzyme *HpaII* and MspI respectively.

**Fig 3.**
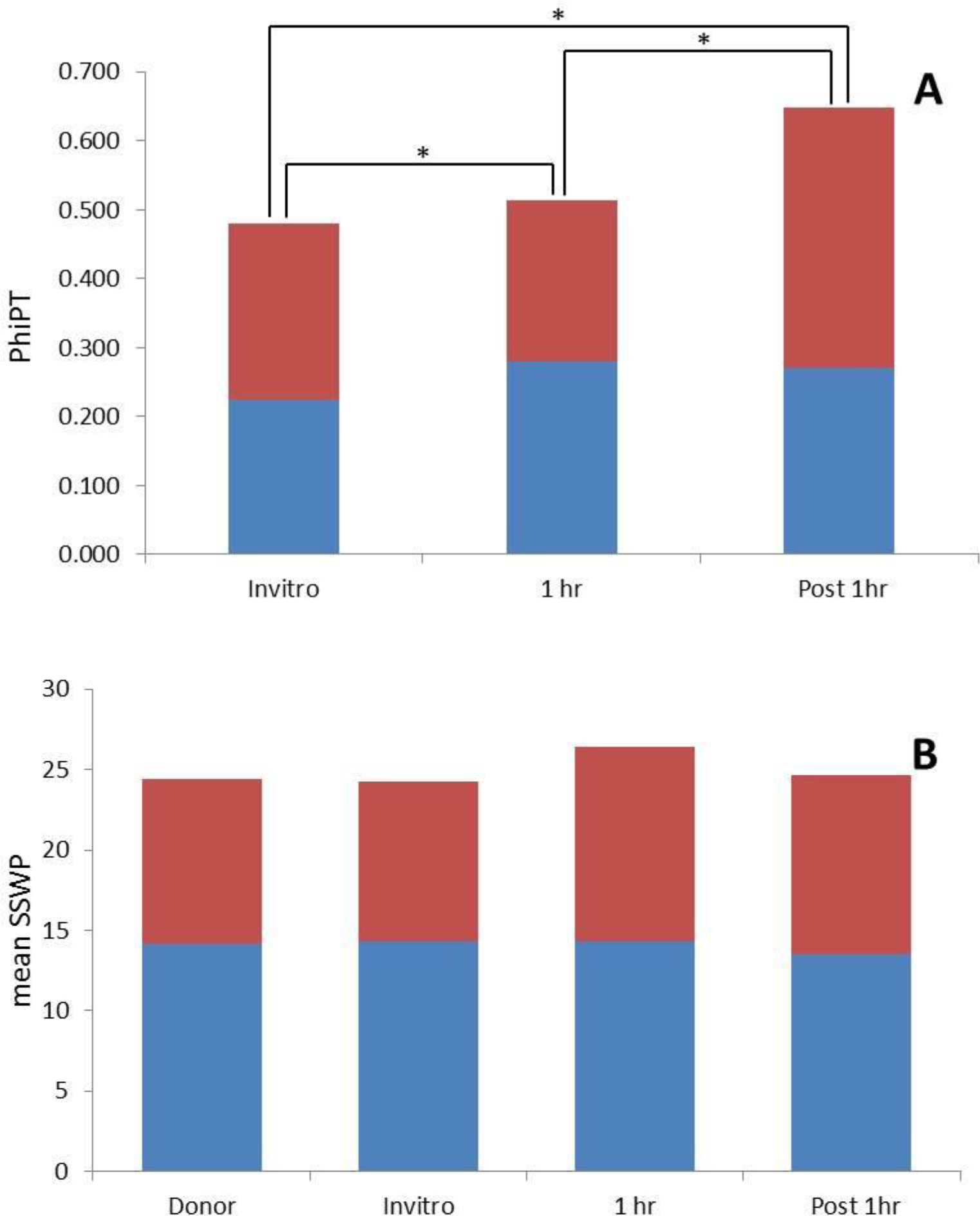
Somatic embryogenesis and cryopreservation induced between epigenetic variability. (A) Bars represent epigenetic distance (PhiPT) between ortet tree samples and each of the *in vitro* treatments (**Invitro**: *In vitro* maintained secondary somatic embryos; **1 h LN**: Secondary somatic embryos recovered from after 1 hour storage in liquid nitrogen and **Post 1 h LN**: tertiary somatic embryos generated from 1 h LN samples) calculated using 10.000 permutations and AMOVA analysis. **(B)** Bars represent the average epigenetic variability (mean sum of squares within population (SSWP)) between samples with the same origin (i.e. Donor, Invitro, 1 h LN and Post 1 h LN). PhiPT and SSWP values were calculated using GenAlex 6.1 software from MSAP profiles generated combining *Hp3/Eco3* and *Hp1/Eco1* selective primer combinations and restriction enzymes *HapII* (Blue) and MspI (red) as frequent cutters. All somatic embryos and ortet tree samples (24 per treatment) were initiated/collected from a single AMAZ 15 cocoa tree from the International Cocoa Quarantine Centre, Reading, UK. * Indicates significantly different (P>0.0002) PhiPT values between treatments.

**Table 4.** Somatic embryogenesis and cryopreservation induced epigenetic differences. Epigenetic distances (PhiPT) between groups (i.e. **Donor:** AMAZ 15 cocoa ortet tree used to regenerate all somatic embryos; **Invitro**: *In vitro* maintained secondary somatic embryos; **1 h LN**: Secondary somatic embryos recovered from after 1 h storage in liquid nitrogen and **Post 1 h LN**: tertiary somatic embryos generated from 1 h LN samples) were calculated using Analysis of Molecular Variance (AMOVA) inferred from the analysis of methylation-sensitive amplified polymorphism (MSAP) assays using methylation sensitive isoschizomers *HpaII* and MspI and primer combinations *Hp1/Eco1* and *Hp3/Eco3.* PhiPT Values are shown below diagonal (-). Probability of having a more extreme PhiPT than the observed values by chance alone based on 10,000 permutations is shown above diagonal.

|  |  | Donor | Invitro | 1 h LN | Post 1h LN |
| --- | --- | --- | --- | --- | --- |
| *HpaII* | Donor | - | 0.0001 | 0.0001 | 0.0001 |
|  | Invitro | 0.2232 | - | 0.0001 | 0.0002 |
|  | 1 h LN | 0.2798 | 0.1401 | - | 0.0001 |
|  | Post 1h LN | 0.2707 | 0.0658 | 0.0778 | - |
| *MspI* | Donor | - | 0.0001 | 0.0001 | 0.0001 |
|  | Invitro | 0.2559 | - | 0.0001 | 0.0001 |
|  | 1 h LN | 0.2340 | 0.0896 | - | 0.0001 |
|  | Post 1h LN | 0.3764 | 0.1206 | 0.1536 | - |

## Discussion

### Somaclonal variation among conserved somatic embryos

Conventional propagation of elite cocoa clones via cuttings has been hampered by costs and undesired phenotypes associated with this type of propagule. Moreover, the long term conservation of cocoa is hindered by high levels of variability in agronomic performance and the poor low temperature storage ability of cocoa seeds [33]. For these reasons considerable effort has been put into the development of *in vitro* propagation and cryopreservation systems (for an extensive review see [34]). However, the passage of cells through *in vitro* culture [11] and cryopreservation [35] may lead to undesirable changes (somaclonal variation) in the regenerants. Although the effect of *in vitro* culture, somatic embryogenesis and cryopreservation of somatic embryos on the morphological and genetic fidelity of cocoa plants has been extensively studied [12,14–16,36], very little is known about the epigenetic modifications introduced by the cryopreservation of cocoa somatic embryos.
Not surprisingly, our results showed an increase in variability in all *in vitro* cultured samples (*in vitro*, 1h LN, Post 1h LN). As we have shown previously, this variability is both genetic and epigenetic in nature [12,14–16,36]. More interestingly, SSE recultured after one hour in liquid nitrogen (i.e. 1h LN and Post-1h LN) showed an increase in genetic/epigenetic distance from the donor samples, and higher levels of within group genetic/epigenetic variability than donor plant samples and thannon-cryopreserved somatic embryos. High frequencies of morphological abnormalities in cryostored and non-cryostored cocoa somatic embryos have been reported [37–39] suggesting that both somatic embryogenesis and cryopreservation can induce somaclonal variability. Evidence from other plant species such as papaya [35] and chrysanthemum [40] suggests that such variation is both genetic and epigenetic (i.e. DNA methylation) in nature. However, our previous work has shown that the genetic variability attributable to the encapsulation-dehydration and vitrification cryopreservation of cocoa SE methodologies used here [29,41] is negligible [16,36]. We, therefore, speculate that this observed increased within group variability is mainly epigenetic in nature, as previously shown in somatic embryo-derived oil palm [42] and cryopreserved hop plants [13]. Moreover, this higher level of within group variability observed in cryopreserved samples (1 h LN) supports the hypothesis that a large fraction of stress (cold, dehydration, etc.) induced epigenetic variation occurs randomly across the genome [43,44].

Interestingly, TSE regenerated from cryopreserved SSE (Post 1h LN) showed intermediate within group genetic/epigenetic variability (mean SSWP*_mspl_* = 264). This would indicate that the cryopreservation induced variability observed in 1h LN samples is partially reversible, supporting the hypothesis of it being epigenetic. Conversely, tertiary somatic embryos in the Post 1h LN group showed a significantly higher PhiPT distance from donor samples than that observed for SSE *(in vitro* and 1h LN samples), suggesting that this extra somatic embryogenesis event does introduce additional genomic variability. Analysis of scanning electron microscopy images have shown that TSE after cryopreservation generally regenerate directly from SSE epidermal cells [16] without the intermediate callus phase which is normally associated with higher levels of somaclonal variation. Nonetheless, epigenetic somaclonal variation is not limited to the passage through a dedifferentiated phase as shown in micropropagated cassava [26]. Furthermore, this increase in variation following another step of somatic embryogenesis is in line with a number of studies on *in vitro* culture variation which report that variation increases with the number of multiplication cycles [3,13]. However, we have also shown that successive rounds of somatic embryogenesis in cocoa reduce the detected levels of genetic variability [16]. It is therefore tempting to speculate that this increase in PhiPT observed using MSAPs is also mainly epigenetic in nature. A factor possibly responsible for this elevated epigenetic variability might be the presence of growth regulators (2,4-D and 6-BA at 1 mg/l and 50 μl /l respectively) used during the induction of TSE, which have been reported to contribute to epi-mutations in tissue culture derived materials [45].

Finally, it is widely accepted that in general the majority of the developmental epigenetic variability occurs randomly across the genome [43,44]. It would therefore be expected that the accumulation of stochastic variation during successive somatic embryogenesis events would increase the observed differences between samples of the same group. If this was the case, then non cryopreserved SSEs should present higher levels of within group variability than donor samples but lower than TSE. However, we observed similar levels of within group variability, measured here as mean SSWP, among all compared groups (donor, *in vitro*, and post 1h LN) suggesting that the somatic embryogenesis-induced variability detected here is not stochastic but probably resulted from a directed adaptation to the successive growing environments to which SEs are exposed.

## Conclusions

The results shown here support previous evidence [16] that the high levels of phenotypic variability observed in cryostored cocoa SE may be symptomatic of epigenetic change. More importantly, our results suggest that this observed variability is not necessarily stochastic in nature, but might be partially a response to the environmental stresses that plant cells are exposed to during *in vitro* culture and cryopreservation. One would expect that the main contributor to this observed variability would be low temperature. In fact RNA directed DNA methylation is upregulated by low temperature [46]. However previous work has shown that low temperature alone does not explain all the variability observed during cryopreservation. For example, the use of the cryoprotectant DMSO, which chelates to nucleic acids, as a cryoprotectant might not only be introducing genetic changes, as shown previously [47] but may be potentially causing numerous DNA methylation changes. What is more, Harding et al. [48] reported that changes on methylation of DNA sequences may be an adaptive response to conditions of high osmotic stress induced by the used of high concentration of sucrose during vitrification. Understanding how these factors affect the fidelity of the regenerant plants will ultimately help in the development of new cryopreservation methodologies with lower levels of phenotypic, genetic and epigenetic variation.

## Supporting Information

**S1 Fig.**
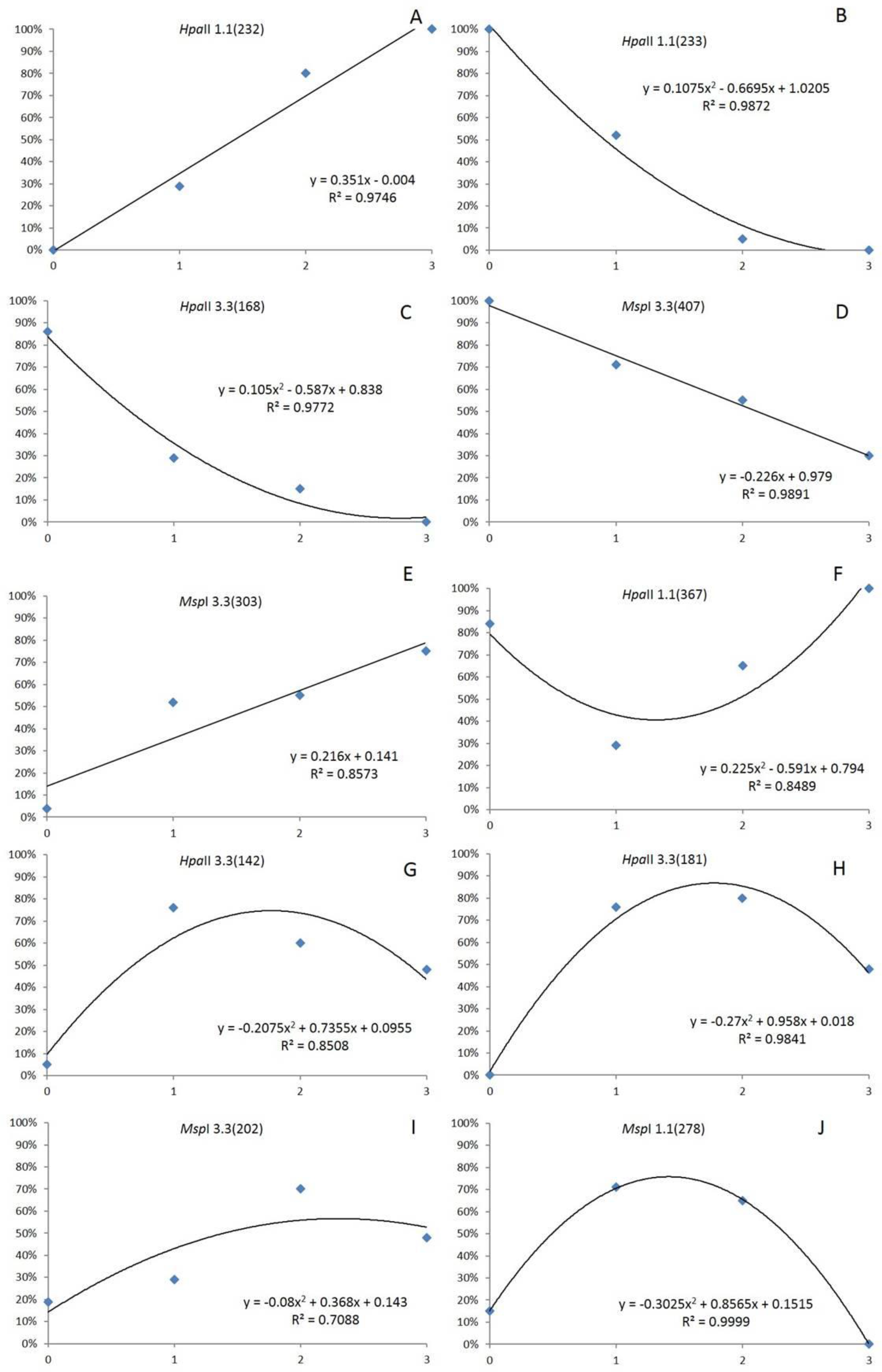
Effect of somatic embryogenesis and cryopreservation on epilocus frequency. Values on the horizontal axis indicated the number of treatments each group has been subjected to (0= Donor Plant, 1= secondary somatic embryos maintained in ED medium, 2= secondary somatic embryos after cryopreservation for 1 h in liquid nitrogen (1h LN) and 3= tertiary somatic embryos generated from 1 h LN samples). Epilocus frequencies were calculated from presence/absence MSAP profiles generated combining *Hp3/Eco3* and *Hp1/Eco1* selective primer combinations and restriction enzymes *HapII* and *MspI* using GenAlex 6.1 software.

